# Reply to: Caution Regarding the Specificities of Pan-Cancer Microbial Structure

**DOI:** 10.1101/2023.02.10.528049

**Authors:** Gregory D. Sepich-Poore, Evguenia Kopylova, Qiyun Zhu, Carolina Carpenter, Serena Fraraccio, Stephen Wandro, Tomasz Kosciolek, Stefan Janssen, Jessica Metcalf, Se Jin Song, Jad Kanbar, Sandrine Miller-Montgomery, Robert Heaton, Rana Mckay, Sandip Pravin Patel, Austin D Swafford, Rob Knight

## Abstract

The cancer microbiome field tremendously accelerated following the release of our manuscript nearly three years ago^1^, including direct validation of our cancer type-specific conclusions in independent, international cohorts^2,3^ and the tumor microbiome’s adoption into the hallmarks of cancer^4^. Disentangling contamination signals from biological signals is an important consideration for this research field. Therefore, despite numerous, high-impact, peer-reviewed research papers that either validated our conclusions or extended them using data we released^2,5–13^, we carefully considered criticism raised by Gihawi *et al*. about potential mishandling of contaminants, batch effects, and machine learning approaches—all of which were central topics in our manuscript. Nonetheless, a close examination of each concern alongside the original manuscript and re-analyses of our published data strongly demonstrates the robustness of the original findings. To remove all doubt, however, we have reproduced all key conclusions from the original manuscript using only overlapping bacterial genera identified in a highly decontaminated, multi-cancer, international cohort (Weizmann Institute of Science, WIS)^2^, with or without batch correction, and with multiclass machine learning analyses to mitigate class imbalances. Our published pan-cancer mycobiome manuscript^3^ also affirms these findings using updated, state-of-the-art methods. We also note that every analysis shown here was possible using public data and code that we had already provided.

## Main

As late as 2015, the tumor microbiome was considered an elusive ‘mirage’^14^, but the view that it was real quickly solidified after the discovery of chemo-degrading bacteria in >75% of pancreatic cancers^15^. Subsequent studies annotated the functional, often immunomodulatory, impacts of these intra-pancreatic bacteria^16,17^ and even fungi^18^, followed by characterization of microbes in non-gastrointestinal cancer types, including lung cancer^19^ and leukemia^20^. Prior to our manuscript, however, multi-cancer microbiome profiling was rare, and the largest attempts excluded ∼85% of The Cancer Genome Atlas (TCGA) patients while lacking systematic decontamination, batch correction, cross-cancer comparisons, or blood-related analyses^21^. In contrast, our paper^1^ analyzed microbial abundances across all 33 TCGA cancer types, with standardized methods for batch correction, *in silico* decontamination, and machine learning (ML) comparisons. These approaches allowed us to conclude that microbial compositions were distinct between and within cancer types, and that trace amounts of their DNA were detectable in human blood and plasma samples, thereby suggesting a novel diagnostic approach^1^. Moreover, these conclusions were later validated and extended by numerous papers^2,5–13^.

In their comments on our study, Gihawi *et al*. begin by stating that most ML models appear to perform no better than using no information and claim “this was not clear in the main text.” However, their purported issue may arise from not considering how confusion matrices are constructed, as well as a failure to consider the relevant text on the website, which clearly states in red text, “Note: A 50% probability cutoff may *not* always be the best choice for class discrimination.” Specifically, by the definition of creating a confusion matrix, a probability cutoff must be chosen to determine which samples are categorized as “positive class” or “negative class” based on the predicted sample probabilities. However, the choice of a probability cutoff is *arbitrary* and varies depending on the goal of the analysis (e.g., higher specificity or sensitivity). The correct, robust choice is to measure areas under the receiver operating characteristic (ROC) and precision-recall (PR) curves. In fact, the ROC and PR curves are drawn by measuring the sensitivity, specificity, and precision at every possible probability cutoff, such that for as many points that exist on the ROC or PR curves (i.e., hundreds to thousands), there exist an equivalent number of confusion matrices, each with their own corresponding accuracy rates and “P-Value [Acc > NIR]” values. Moreover, this fact is why probability cutoff optimization approaches exist in the first place, including for imbalanced classes (see, for example, pages 423-425 of Kuhn & Johnson^22^).

Because ∼600 ML models are provided on the website, a default 50% probability cutoff was used for the confusion matrices, while clearly noting that other cutoffs may be better in red text (shown above) and clearly showing the full ROC and PR curves. Thus, the question that Gihawi *et al*. should be asking is *not* if the accuracy p-value of an arbitrarily-chosen confusion matrix is significant, but rather if the areas under the ROC (AUROC) and PR (AUPR) curves are greater than their null values, which is 0.5 for AUROC and the positive class prevalence for AUPR. Indeed, *all* the ML models that Gihawi *et al*. claim are concerning substantially exceed these null AUROC and AUPR values, meaning they substantially outperform no-information models.

The precision-recall (PR) curves comprise another fundamental statistical concept that is not discussed by Gihawi *et al*., even though it resolves key elements of their concern regarding ML class imbalances. Specifically, Gihawi *et al*. state “the poor performance of the models may in part be due to the major imbalance in class size in the datasets…” but fail to understand that the formulas for neither precision nor recall use true negatives in their calculations (see, for example, page 74 of Brownlee^23^). This means that AUPR is *not* affected by class imbalance, including during one-versus-all-other ML modeling containing mostly negative class samples. Thus, because all of the cross-cancer ML models have AUPR values that dramatically exceed their null values, it is categorically false to state that the models have “poor performance[s]” due to the class imbalances—they in fact do not. For example, in the adrenocortical carcinoma ML model that Gihawi *et al*. highlight, the null AUPR is ∼0.55% but the measured AUPR is ∼89.55%.

In another section of comments, Gihawi *et al*. raise concerns about how nonsensical genera appear to be informative of tumor type. However, the only feature set that Gihawi *et al*. discuss in this context is the “most stringent decontaminated” dataset, which is the only dataset that removes all typically-expected commensals (see Extended Data Fig. 6c-f in original manuscript) and which was included as a technical control to address reviewer concerns. In fact, when running SourceTracker2^24^ on adjacent normal colon TCGA samples against Human Microbiome Project samples, we “found that successively stringent decontamination improved recognition of concomitant tissues *before they became unrecognizable*”^1^. Furthermore, after the most stringent decontamination, normal colon samples no longer looked like stool (see Extended Data Fig. 6f in original manuscript). Correspondingly, this result led us to conclude, “These results suggest that stringent filtering may be desirable in certain comparisons, but a universal approach to decontamination *may preclude biologically informative results*.”^1^ Indeed, this is why we included *four* decontaminated versions of the data for the reader (e.g., see Figs. 3a-c in the original manuscript), because a gold standard for *in silico* decontamination does not yet exist. Moreover, intersecting the publicly-available, WIS-identified^2^ bacterial genera against each of the datasets we released clearly shows that the most stringent decontaminated data contains <5% overlap, whereas all other datasets have >52% overlap (**Fig. 1a**).

**Fig. 1.**
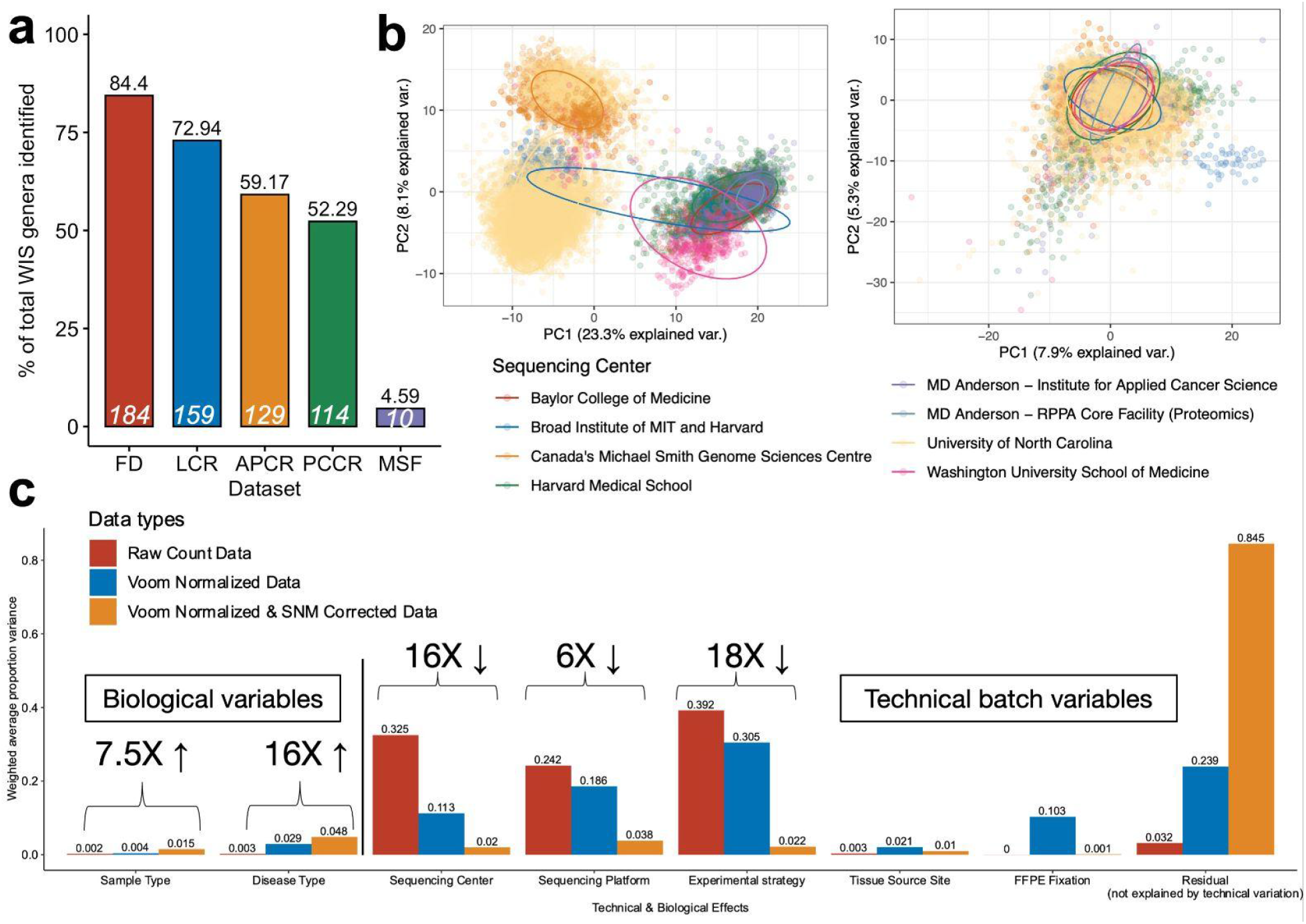
Approach for re-evaluating the TCGA cancer microbiome analysis using WIS-overlapping features. **a**, Bar plot depicting the bacterial genera overlap between those identified by Nejman *et al*. (WIS) and those in the original Kraken data published by Poore *et al*., including the percentage of WIS-genera covered in each released dataset (black labels) and the absolute number of genera covered (white labels). The total number of WIS bacterial genera was 218. FD, full data; LCR, likely contaminants removed by sequencing center; APCR, all putative contaminants removed by sequencing center; PCCR, plate–center contaminants removed; MSF, most stringent filtering by sequencing center. **b**, Principal components analysis (PCA) of Voom-normalized data before batch correction (left) and Voom-SNM data post batch correction (right) with cancer microbiome samples coloured by sequencing center. **c**, Principal variance components analysis of raw taxonomical count data, Voom-normalized data, and Voom-SNM data.

An additional conceptual flaw in the critique is conflating differential abundance with feature importance scores, which provide neither directionality (i.e., the most important feature can be associated with the negative class) nor significance (i.e., there is no guarantee that any ranked feature is significantly over-or under-abundant in either class). As with the confusion matrix cutoff concern above, we correctly stated this on the original website in red text:

> “Moreover, a high feature importance score for a given taxon does *not* guarantee or imply an overabundance of that taxon for this comparison. A high feature importance score only means that the taxon was important for making predictions; whether the taxon is less or more abundant in certain samples requires statistical testing.”

Gihawi *et al*. claim overwhelming contamination in our data, while neglecting all of the positive control analyses we conducted in the original manuscript (Fig. 2a-g in original text). For viruses, these analyses included verifying significant overabundance of HPV in cervical cancers and head and neck cancers that were clinically positive for HPV (Figs. 2d-e in original text); demonstrating that *Lymphocryptovirus* was significantly overabundant in the corresponding subtype of stomach cancer^25^ (Fig. 2g in original text); and showing that hepatitis B virus (HBV) was enriched in concomitantly HBV-infected liver cancers (Fig. 2f in original text). For bacteria, these analyses included verifying tumor tissue-specific *Fusobacterium* overabundance in eight gastrointestinal cancers (Figs. 2b-c in original text); verifying the presence, but not differential abundance^25^, of *Helicobacter* in stomach cancer tumor vs. normal tissue; and revealing that normal colon samples indeed looked like stool via source tracking (Fig. 2a in original text). All of these analyses demonstrated that expected taxa were found in both anticipated presence/absence and relative abundances across different sample types. In fact, another paper by Gihawi *et al*.^26^ also tested a subset of the same positive controls using tumor sequencing data and found the same results—*Alphapapillomavirus* in cervical cancer, *Helicobacter* in stomach cancer, *Lymphocryptovirus* in stomach cancer—and concluded that these validated their dataset and analyses.

**Fig. 2.**
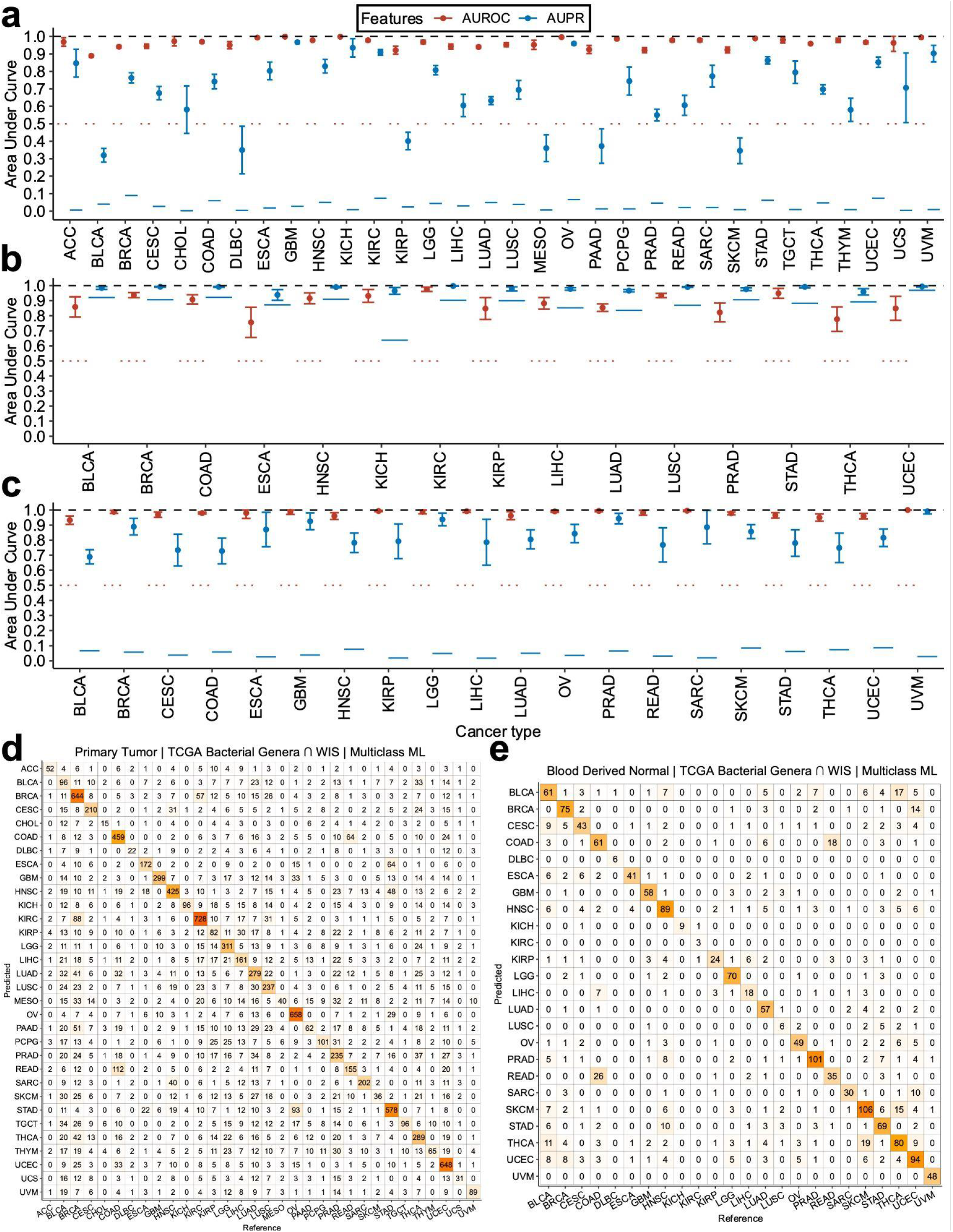
Classifier performance for cancer discrimination using WIS-overlapping bacterial genera between and among cancer types using tissue and blood-derived microbial nucleic acids. **a**, One-cancer-type-versus-all-others predictions using WIS-overlapping, batch-corrected, TCGA primary tumor data, with gradient boosting machines and 10-fold cross validation. **b**, Primary tumor tissue versus adjacent tissue normals per cancer type using WIS-overlapping, batch-corrected, TCGA tissue data, with gradient boosting machines and 10-fold cross validation. **c**, One-cancer-type-versus-all-others predictions using WIS-overlapping, batch-corrected, TCGA blood data, with gradient boosting machines and 10-fold cross validation. **d**, Multiclass machine learning amongst all primary tumor types simultaneously in TCGA using WIS-overlapping, batch-corrected, TCGA primary tumor data, with gradient boosting machines and 10-fold cross validation. **e**, Multiclass machine learning amongst all cancer types simultaneously in TCGA using WIS-overlapping, batch-corrected, TCGA blood data, with gradient boosting machines and 10-fold cross validation. **a, b, c**, AUROC and AUPR measured on independent holdout folds (ten-fold cross-validation (CV)) to estimate averages (dots) and 95% confidence intervals (brackets). Red horizontal dotted lines under AUROC denote null values. Blue solid horizontal lines under AUPR denote null values, which equates the prevalence of the positive class (each cancer type) among the full set of all cancer types.

**Fig. 3.**
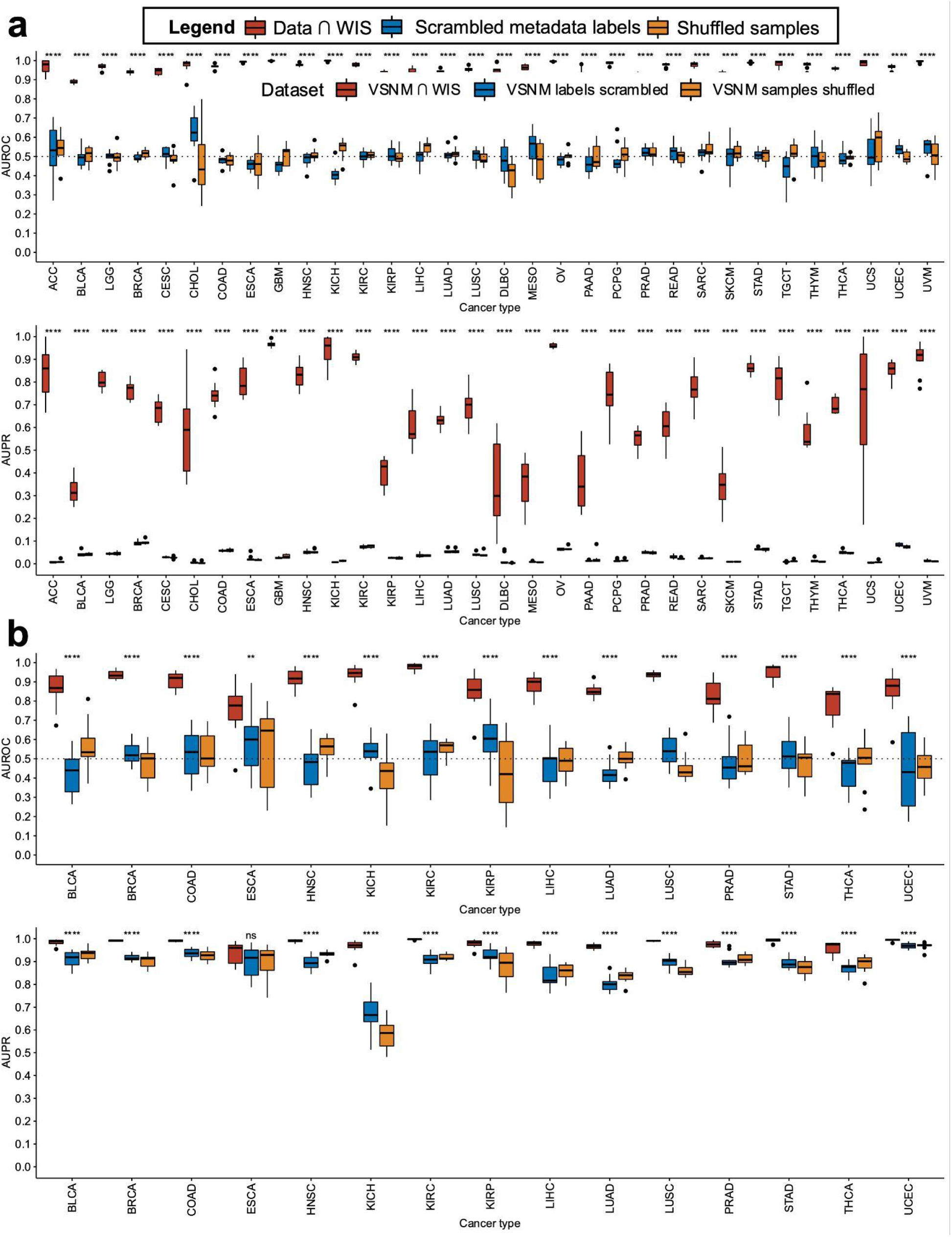
Control analyses to verify TCGA tissue-based classifier performances by comparing against ML models built with scrambled metadata or shuffled samples. **a**, Primary tumor-based ML models (cf. **Fig. 2a**) were repeated using scrambled metadata labels or shuffled count data and compared to the original performances using biological samples with correct labels. **b**, Primary tumor vs. adjacent tissue normal-based ML models (cf. **Fig. 2b**) were repeated using scrambled metadata labels or shuffled count data and compared to the original performances using biological samples with correct labels. **a, b**, For hypothesis testing, biological data and scrambled/shuffled controls were aggregated into two separate groups, and two-sided Wilcox tests were applied per cancer type per performance metric (AUROC or AUPR). *: p≤0.05; **: p≤0.01; ***: p≤0.001; ****: p≤0.0001.

In their next section, Gihawi *et al*. make a case that the observed nonsensical genera are likely traceable to human read misclassification. Although we endorse bioinformatic protocols that extensively remove human reads from metagenomic data, the papers that Gihawi *et al*. cite^26,27^ as evidence for additional host depletion steps were published *months after* our manuscript was already submitted (June 7, 2019; stated on the published manuscript) or published online (March 11, 2020). To critique that our study should have implemented bioinformatic protocols or considered benchmarking that were only available *after* our manuscript was submitted or published is not reasonable. Additionally, within our manuscript, we performed direct genome alignments of Kraken-positive, genus-level microbial reads from a subset of four cancer types—cervical squamous cell carcinoma (CESC), stomach adenocarcinoma (STAD), lung adenocarcinoma (LUAD), and ovarian serous cystadenocarcinoma (OV)—finding “a low estimated false-positive rate of 1.09% (Supplementary Table 3)” (quoted from the original manuscript). Moreover, our more recent pan-cancer mycobiome manuscript^3^ systematically implements these additional, extensive bioinformatic host filtering methods, and reached the same cancer type-specific conclusions as our original manuscript^1^.

In subsequent comments, Gihawi *et al*. raise concerns about the batch effect method employed in our study, namely Voom^28^, which was used to transform heteroscedastic microbial data into pseudo-normally distributed data. Specifically, Gihawi *et al*. argue that Voom’s pseudocount for zero-valued features is inappropriate and “raises prominent concerns at the level of individual taxa.” However, Voom was precisely designed to transform heteroscedastic data into pseudo-normally distributed data,^28^ and has been cited by >4000 research papers to date, including the majority of papers that apply it for RNA-Seq differential expression analyses on individual genes. In each of these applications of Voom, the algorithm adds a pseudocount to zero-valued features, which is maintained during subsequent differential expression modeling and calculation of log-fold changes on individual genes. To reject Voom’s methodology would require discarding a substantial portion of the literature, and Gihawi *et al*. fail to offer a better alternative. We also note that other published approaches for microbial differential abundance, such as ANCOM-II, also apply pseudocounts prior to log transformation steps^29^. Moreover, in our application, Voom preserved expected sample type and clinically-related differences (e.g., tumor vs. normal tissue, HPV-positive vs. HPV-negative tumors) across thousands of patient samples in our positive control analyses (Fig. 2 in original text). Furthermore, the magnitude of expected and observed differences between sample types and clinically-related differences were substantially larger in comparison to the concern by Gihawi *et al*. of Voom-induced statistical noise. For example, HPV-positive cervical tumors had a median Voom-SNM value of 8.69, whereas HPV-negative tumors had a median value of 0.64 (Fig. 2d source data, original text), or a percent change of 1257.81%; in contrast, the *Hepandenosovirus* shown by Gihawi *et al*. in Figure 1 has values varying between 3.078 and 3.084, or a maximum percent change of just 0.19%. Nonetheless, we show below (**Figs. 5-10**) and in our pan-cancer mycobiome manuscript^3^ that it is possible to skip Voom altogether, using only the raw microbial counts, and reach the same conclusions as the original manuscript, so even if this concern applied in principle it does not affect the main results.

Throughout their piece, Gihawi *et al*. cite Whalen *et al*.^30^ as evidence that five key ML mistakes were made in our manuscript. However, in every instance, the underlying Whalen principle is either misapplied or already satisfied by analyses included in our manuscript. For instance, Gihawi *et al*. cite “Whalen V: unbalanced classes” in their critique about applying upsampling in our manuscript to correct for imbalanced classes during ML. But Whalen *et al*. actually *recommended* using upsampling as a way to address class imbalance:

> “Class imbalance is addressed with a range of strategies […] There are three basic strategies: *oversampling the minority class*, undersampling the majority class and weighting examples […] Each approach makes different trade-offs: *oversampling retains all data but increases computation time*, undersampling decreases computation time but discards some data and sample weighting retains all data but requires determination of the optimal weights.”^30^

In our analyses, we chose to up/oversample because it did not discard any of the data although it did indeed increase the computational cost substantially. Therefore, our procedure is exactly as recommended by Whalen *et al*. We also note that Whalen *et al*. explicitly discussed the use of AUPR in cases of class imbalance, recommending exactly the procedure we used:

> “Alternatively, auPR compares recall with precision, which is one minus the false discovery rate and does not depend on the number of negatives. When false positives dominate true positives, auPR will be low. Thus, for imbalanced problems where the positive minority class is of primary interest, auPR is generally preferred.”^30^

As another example, in their discussion about Voom-related batch correction, Gihawi *et al*. cite “Whalen IV - leaky preprocessing,” asserting that the batch correction of the dataset prior to ML biased the results. However, we addressed this claim in Extended Data Fig. 3a in the original text^1^, which separated the raw TCGA data into two separate halves, performed batch correction on them independently, trained ML models on each half independently, and then cross-tested, showing the same conclusion (that microbiomes are cancer type-specific). Similarly, when Gihawi *et al*. also state that the batch correction by sequencing center was an example of “Whalen III: confounding,” they ignore Extended Data Figs. 3f-i in the original text^1^, which show how our ML performances were significantly correlated between samples subset to individual sequencing centers and the full dataset.

Similarly, the last two Whalen principles that are raised by Gihawi *et al*. as apparent deficiencies are misapplied. For instance, Gihawi *et al*. assert that the inclusion of bacteriophages violates “Whalen II: dependent examples” since they require a bacterial host in one or more anatomical locations; however, what Whalen *et al*. actually discuss is whether independence between *samples* exists, not features^30^. Because the presence/absence of a bacteriophage in one patient (e.g., with liver cancer) is completely independent from presence/absence in another patient (e.g., with lung cancer), there is no relevant inter-dependence in the data structure we presented. Additionally, when Gihawi *et al*. state, “microbiome data is dynamic[21] (Whalen I: distributional differences),” they suggest that the time-varying qualities of the microbiome preclude modeling. However, Whalen *et al*. actually discuss time-series analyses as a potential issue of ‘Whalen II: dependent examples’—not ‘Whalen I’—since patient samples in time-series data are dependent on each other. TCGA focused on collecting cross-sectional samples from treatment-naive patients and not on time-series collection, making the critique of dynamic microbiomes inapplicable. Collectively, Gihawi *et al*. misapplied each of the Whalen principles and did not consider multiple analyses that specifically address such points.

We agree with Gihawi *et al*. that following “the RIDE criteria set out by Eisenhofer *et al*. (also authored by Knight) as closely as possible” is critical, when possible, for tumor microbiome research. However, in the case of our large-scale analysis of TCGA, three of the four RIDE criteria (‘-IDE’) requiring experimental contamination controls were not applicable since TCGA had none, which we clearly communicated in the original Methods: “As TCGA did not include any negative blank reagent tubes during sample processing…”^1^ Moreover, the sole remaining criterion (‘R-’) that does not rely on experimental controls—to “report the experimental design and approaches used to reduce and assess the contributions of contamination”^31^—was indeed directly diagrammed in Extended Data Fig. 6a in the original text^1^. All of these diagrammed methods attempted to infer the amount and types of contaminating taxa using state-of-the-art in silico methods, which follow the spirit of the RIDE guidelines^31^. Moreover, in our validation experiment with new samples (Fig. 4 in the original text), we indeed implemented both the RIDE guidelines^31^ and KatharoSeq protocol^32^, including negative and serially titrated positive controls on a single plate for sequencing. Thus, when there was the opportunity to use them, we indeed did.

**Fig. 4.**
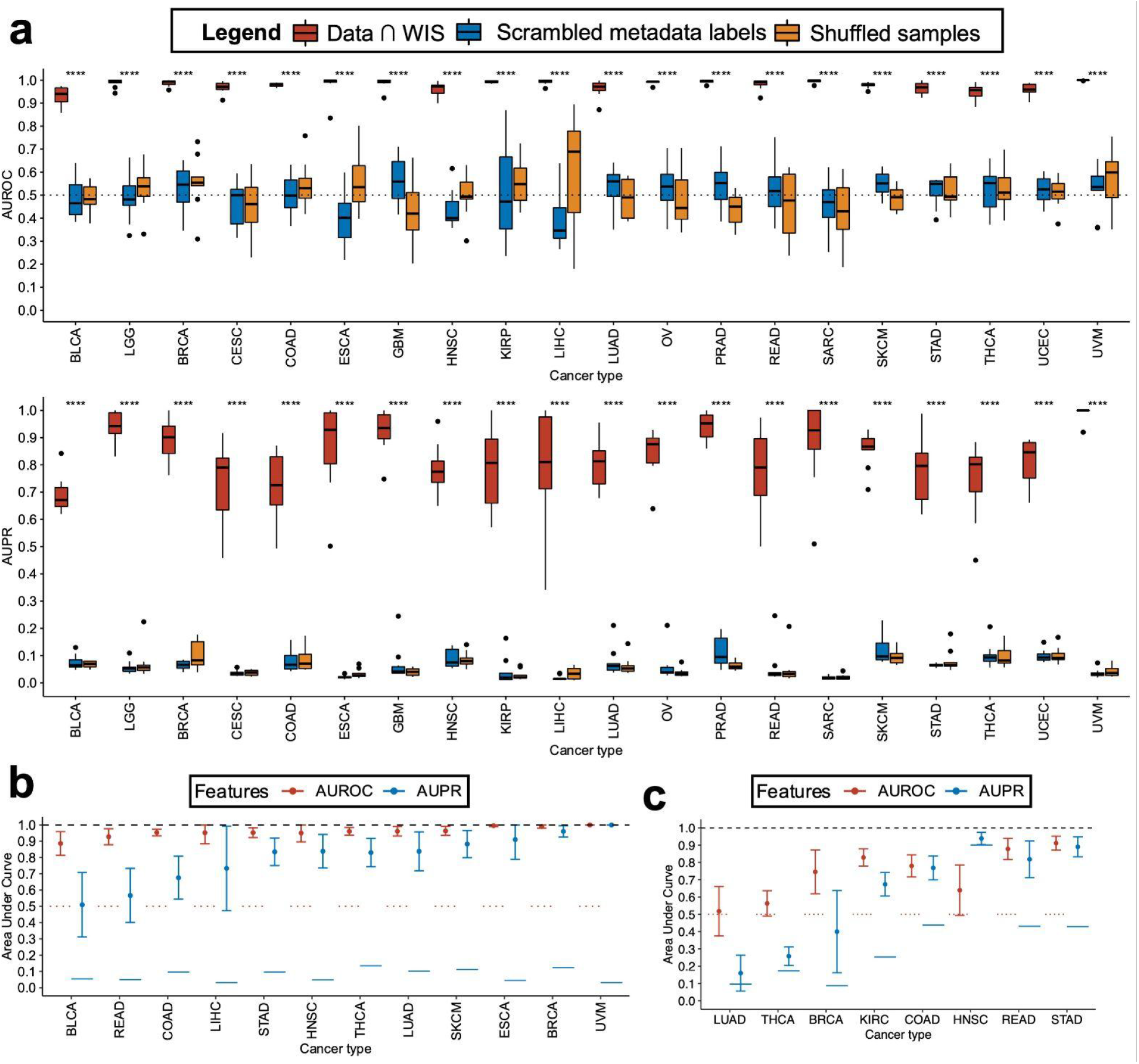
Control analyses to verify TCGA blood-based classifier performances and further ML using blood samples from low-stage cancers or comparing primary tumors from low and high clinical stages. **a**, Blood-based ML models (cf. **Fig. 2c**) were repeated using scrambled metadata labels or shuffled count data and compared to the original performances using biological samples with correct labels. For hypothesis testing, biological data and scrambled/shuffled controls were aggregated into two separate groups, and two-sided Wilcox tests were applied per cancer type per performance metric (AUROC or AUPR). *: p≤0.05; **: p≤0.01; ***: p≤0.001; ****: p≤0.0001. **b**, One-cancer-type-versus-all-others predictions using WIS-overlapping, batch-corrected, TCGA blood data from patients having stage Ia-IIc cancer, with gradient boosting machines and 10-fold cross validation. **c**, Individual cancer predictions using WIS-overlapping, batch-corrected, TCGA primary tumor data to distinguish stage I versus stage IV tumors. **b, c**, AUROC and AUPR measured on independent holdout folds (ten-fold cross-validation (CV)) to estimate averages (dots) and 95% confidence intervals (brackets). Red horizontal dotted lines under AUROC denote null values. Blue solid horizontal lines under AUPR denote null values, which equates the prevalence of the positive class (each cancer type) among the full set of all cancer types.

**Fig. 5.**
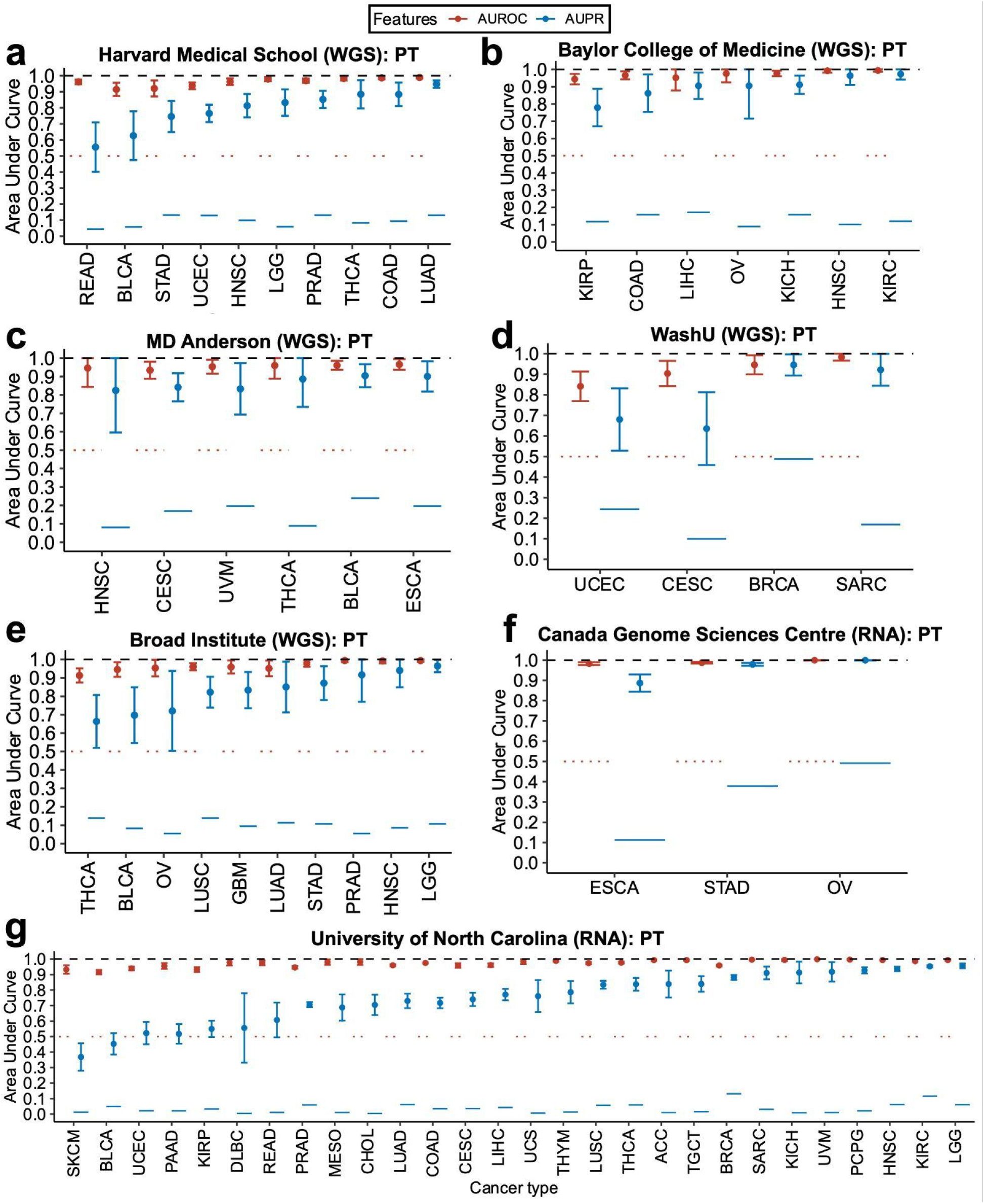
Classifier performance using subset raw data with WIS-overlapping bacterial genera for cancer discrimination using primary tumor-derived microbial nucleic acids. Raw count data using WIS-overlapping bacterial genera were subset to a single sequencing center, experimental strategy (WGS or RNA-Seq), and sequencing platform (Illumina HiSeq) prior to evaluating ten-fold cross-validation gradient boosting machine learning models. Notably, 6 of 7 sequencing centers only used one type of experimental strategy (WGS or RNA-Seq); the exception was the Broad Institute, which performed RNA-Seq on glioblastoma tumors, but since it was the only cancer type to have RNA from Broad, it could not be compared in one-cancer-type-versus-all-other predictions. Predictions were made on each of the ten holdout folds to generate average and 95% confidence intervals of discriminatory performance, as measured by AUROC and AUPR. A minimum of 20 samples were required in any comparison to be tested. **a**, One-cancer-type-versus-all-others predictions among WGS primary tumor samples from Harvard Medical School. **b**, One-cancer-type-versus-all-others predictions among WGS primary tumor samples from Baylor College of Medicine. **c**, One-cancer-type-versus-all-others predictions among WGS primary tumor samples from MD Anderson. **d**, One-cancer-type-versus-all-others predictions among WGS primary tumor samples from Washington University. **e**, One-cancer-type-versus-all-others predictions among WGS primary tumor samples from the Broad Institute. **f**, One-cancer-type-versus-all-others predictions among RNA-Seq primary tumor samples from Canada’s Michael Smith Genome Sciences Centre. **g**, One-cancer-type-versus-all-others predictions among RNA-Seq primary tumor samples from the University of North Carolina. **a, b, c, d, e, f, g**, AUROC and AUPR measured on independent holdout folds (ten-fold cross-validation (CV)) to estimate averages (dots) and 95% confidence intervals (brackets). Red horizontal dotted lines under AUROC denote null values. Blue solid horizontal lines under AUPR denote null values, which equates the prevalence of the positive class (each cancer type) among the full set of all cancer types within a sequencing center-experimental strategy-platform subset. PT, primary tumor.

**Fig. 6.**
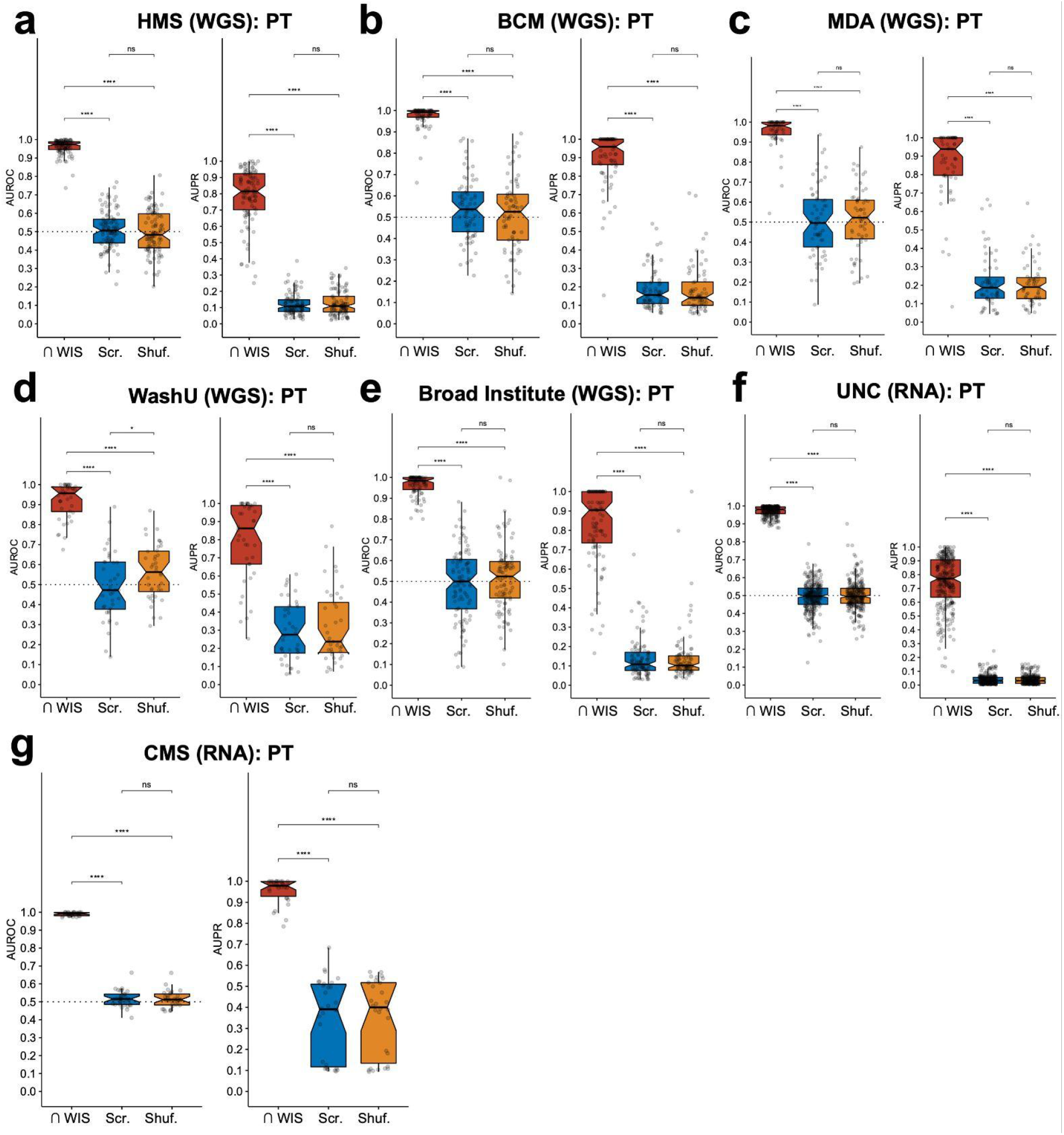
Raw data control analyses to verify TCGA primary tumor classifier performances. Raw count data using WIS-overlapping bacterial genera were subset to a single sequencing center, experimental strategy (WGS or RNA-Seq), and sequencing platform (Illumina HiSeq) prior to evaluating ten-fold cross-validation gradient boosting machine learning models. Performances on these raw data subsets (cf. **Fig. 5**) were then compared to equivalent ML models built with scrambled metadata or shuffled samples. **a**, Comparison of primary tumor-based ML models built on raw count data versus equivalent ones built using scrambled metadata labels or shuffled count data using WGS samples from Harvard Medical School (HMS). **b**, Comparison of primary tumor-based ML models built on raw count data versus equivalent ones built using scrambled metadata labels or shuffled count data using WGS samples from Baylor College of Medicine (BCM). **c**, Comparison of primary tumor-based ML models built on raw count data versus equivalent ones built using scrambled metadata labels or shuffled count data using WGS samples from MD Anderson (MDA). **d**, Comparison of primary tumor-based ML models built on raw count data versus equivalent ones built using scrambled metadata labels or shuffled count data using WGS samples from Washington University (WashU). **e**, Comparison of primary tumor-based ML models built on raw count data versus equivalent ones built using scrambled metadata labels or shuffled count data using WGS samples from the Broad Institute. **f**, Comparison of primary tumor-based ML models built on raw count data versus equivalent ones built using scrambled metadata labels or shuffled count data using RNA-Seq samples from the University of North Carolina (UNC). **g**, Comparison of primary tumor-based ML models built on raw count data versus equivalent ones built using scrambled metadata labels or shuffled count data using RNA-Seq samples from Canada’s Michael Smith Genome Sciences Centre (CMS). **a, b, c, d, e, f, g**, Pairwise two-sided Wilcox tests using performances from all ML folds, corrected for multiple hypothesis testing using the Benjamini-Hochberg method, are shown. ns: not significant (q>0.05); *: q≤0.05; **: q≤0.01; ***: q≤0.001; ****: q≤0.0001. PT, primary tumor.

**Fig. 7.**
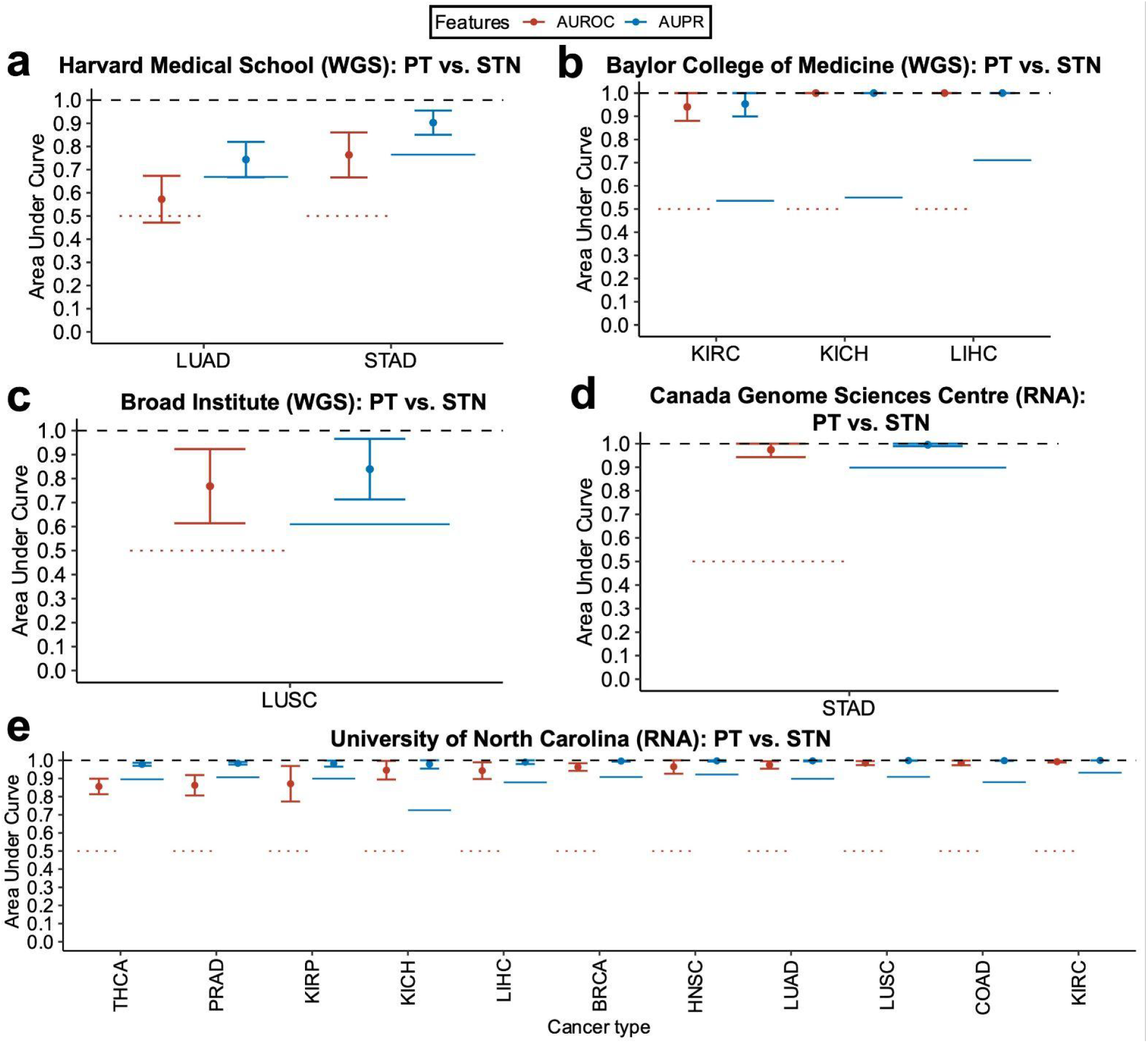
Classifier performance using subset raw data with WIS-overlapping bacterial genera for primary tumor versus adjacent tissue normal discrimination. Raw count data using WIS-overlapping bacterial genera were subset to a single sequencing center, experimental strategy (WGS or RNA-Seq), and sequencing platform (Illumina HiSeq) prior to evaluating ten-fold cross-validation gradient boosting machine learning models. Predictions were made on each of the ten holdout folds to generate average and 95% confidence intervals of discriminatory performance, as measured by AUROC and AUPR. A minimum of 20 samples were required in any comparison to be tested, and neither MD Anderson nor Washington University had sufficient tumor versus normal samples in this setting to evaluate. **a**, Primary tumor vs. adjacent normal using WGS tissue samples from Harvard Medical School. **b**, Primary tumor vs. adjacent normal using WGS tissue samples from Baylor College of Medicine. **c**, Primary tumor vs. adjacent normal using WGS tissue samples from the Broad Institute. **d**, Primary tumor vs. adjacent normal using RNA-Seq tissue samples from Canada’s Michael Smith Genome Sciences Centre. **e**, Primary tumor vs. adjacent normal using RNA-Seq tissue samples from the University of North Carolina. **a, b, c, d, e**, AUROC and AUPR measured on independent holdout folds (ten-fold cross-validation (CV)) to estimate averages (dots) and 95% confidence intervals (brackets). Red horizontal dotted lines under AUROC denote null values. Blue solid horizontal lines under AUPR denote null values, which equates the prevalence of the positive class (each cancer type) among the full set of all cancer types within a sequencing center-experimental strategy-platform subset. PT, primary tumor; STN, solid tissue normal.

**Fig. 8.**
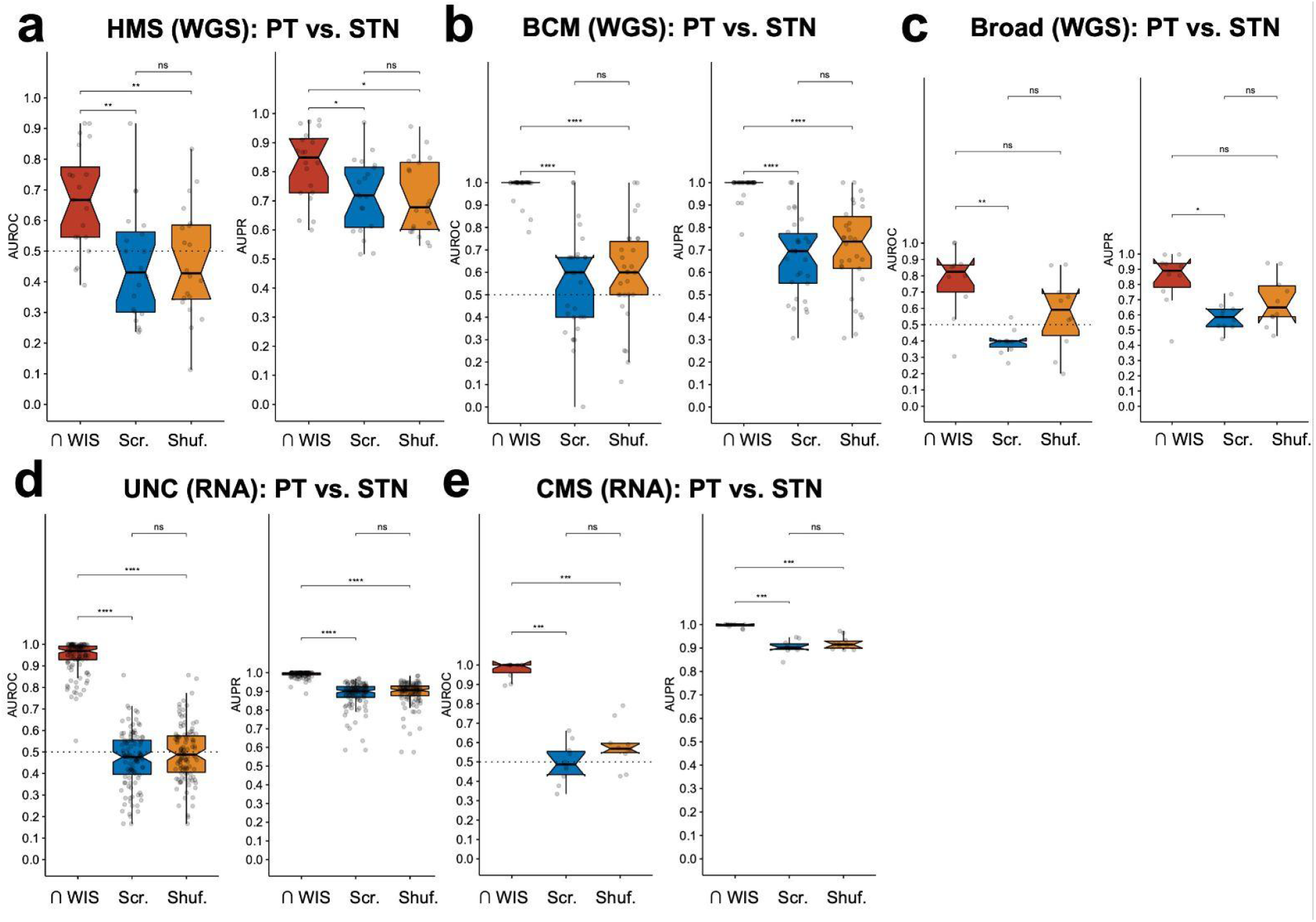
Raw data control analyses to verify TCGA primary tumor vs. adjacent normal classifier performances. Raw count data using WIS-overlapping bacterial genera were subset to a single sequencing center, experimental strategy (WGS or RNA-Seq), and sequencing platform (Illumina HiSeq) prior to evaluating ten-fold cross-validation gradient boosting machine learning models. Tumor vs. adjacent normal performances on these raw data subsets (cf. **Fig. 7**) were then compared to equivalent ML models built with scrambled metadata or shuffled samples. **a**, Comparison of primary tumor vs. adjacent normal ML models built on raw count data versus equivalent models built using scrambled metadata labels or shuffled count data using WGS samples from Harvard Medical School (HMS). **b**, Comparison of primary tumor vs. adjacent normal ML models built on raw count data versus equivalent models built using scrambled metadata labels or shuffled count data using WGS samples from Baylor College of Medicine (BCM). **c**, Comparison of primary tumor vs. adjacent normal ML models built on raw count data versus equivalent models built using scrambled metadata labels or shuffled count data using WGS samples from the Broad Institute (Broad). **d**, Comparison of primary tumor vs. adjacent normal ML models built on raw count data versus equivalent models built using scrambled metadata labels or shuffled count data using RNA-Seq samples from the University of North Carolina (UNC). **e**, Comparison of primary tumor vs. adjacent normal ML models built on raw count data versus equivalent models built using scrambled metadata labels or shuffled count data using RNA-Seq samples from Canada’s Michael Smith Genome Sciences Centre (CMS). **a, b, c, d, e**, Pairwise two-sided Wilcox tests using performances from all ML folds, corrected for multiple hypothesis testing using the Benjamini-Hochberg method, are shown. ns: not significant (q>0.05); *: q≤0.05; **: q≤0.01; ***: q≤0.001; ****: q≤0.0001. PT, primary tumor; STN, solid tissue normal.

**Fig. 9.**
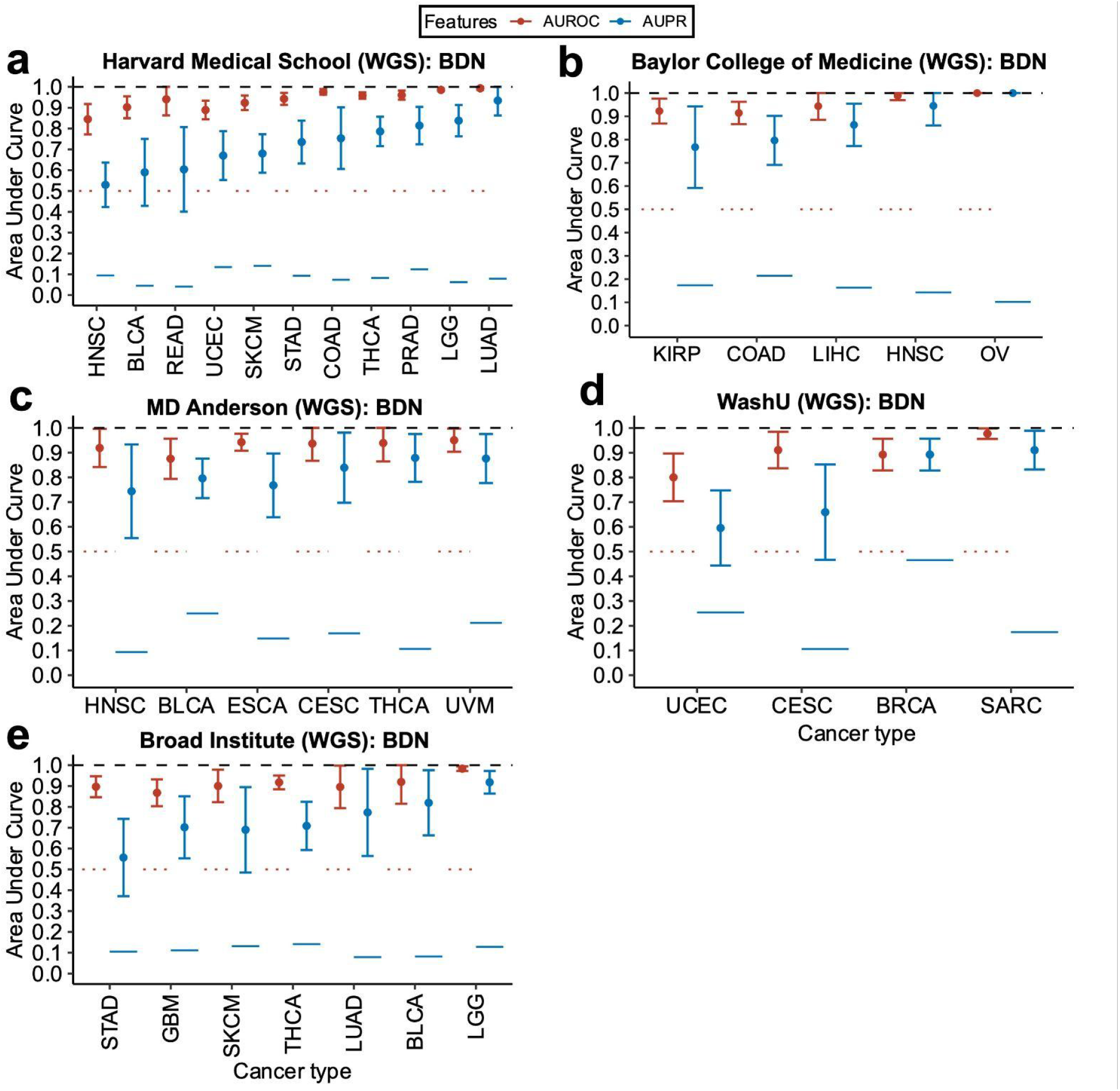
Classifier performance using subset raw data with WIS-overlapping bacterial genera for cancer discrimination using blood-derived microbial nucleic acids. Raw count data using WIS-overlapping bacterial genera were subset to a single sequencing center, experimental strategy (WGS or RNA-Seq), and sequencing platform (Illumina HiSeq) prior to evaluating ten-fold cross-validation gradient boosting machine learning models. All blood derived normal (BDN) samples were WGS. Predictions were made on each of the ten holdout folds to generate average and 95% confidence intervals of discriminatory performance, as measured by AUROC and AUPR. A minimum of 20 samples were required in any comparison to be tested. **a**, One-cancer-type-versus-all-others predictions among WGS blood samples from Harvard Medical School. **b**, One-cancer-type-versus-all-others predictions among WGS blood samples from Baylor College of Medicine. **c**, One-cancer-type-versus-all-others predictions among WGS blood samples from MD Anderson. **d**, One-cancer-type-versus-all-others predictions among WGS blood samples from Washington University (WashU). **e**, One-cancer-type-versus-all-others predictions among WGS blood samples from the Broad Institute. **a, b, c, d, e**, AUROC and AUPR measured on independent holdout folds (ten-fold cross-validation (CV)) to estimate averages (dots) and 95% confidence intervals (brackets). Red horizontal dotted lines under AUROC denote null values. Blue solid horizontal lines under AUPR denote null values, which equates the prevalence of the positive class (each cancer type) among the full set of all cancer types within a sequencing center-experimental strategy-platform subset. BDN, blood derived normal.

**Fig. 10.**
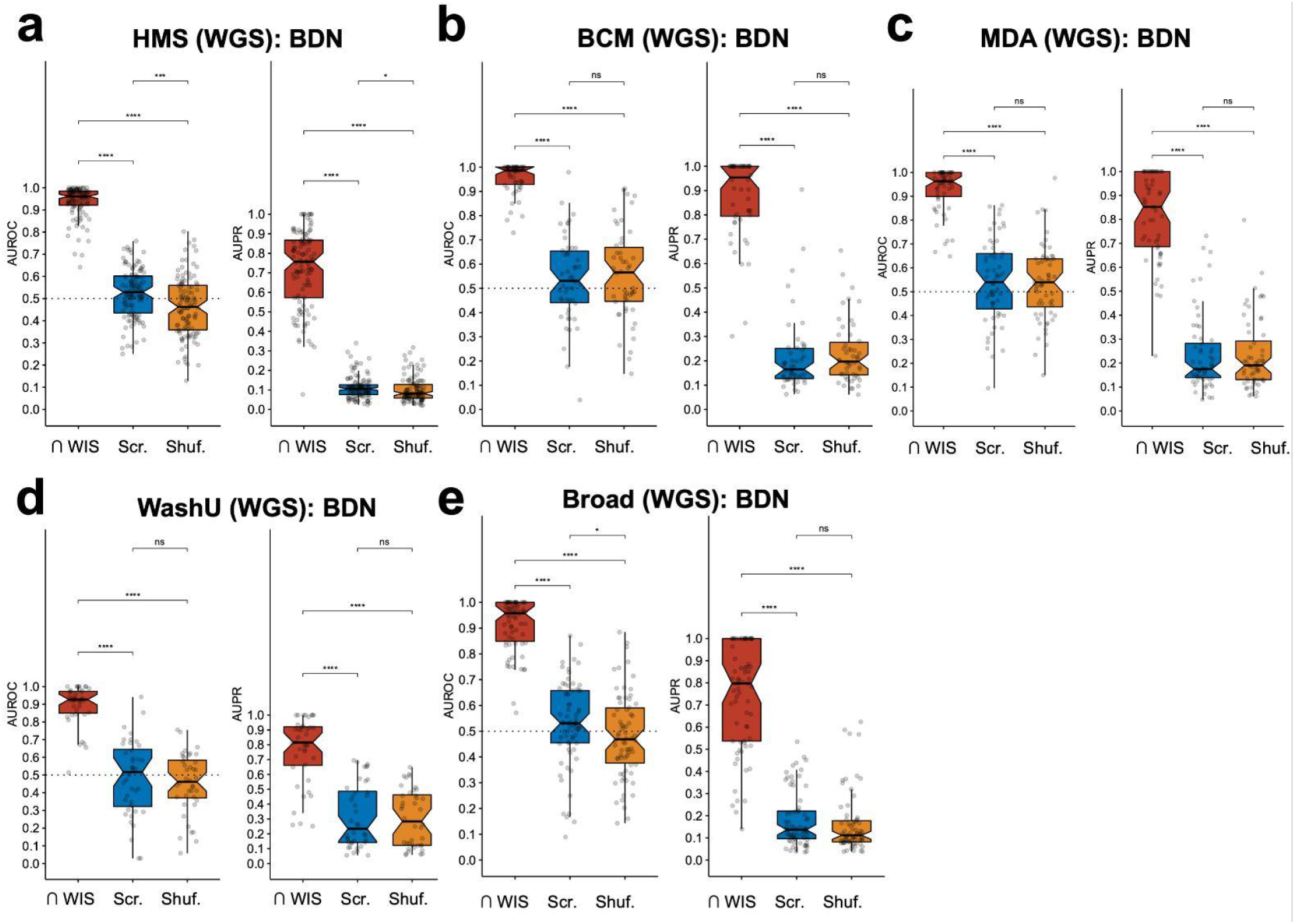
Raw data control analyses to verify TCGA blood classifier performances. Raw count data using WIS-overlapping bacterial genera were subset to a single sequencing center, experimental strategy (all blood samples were WGS), and sequencing platform (Illumina HiSeq) prior to evaluating ten-fold cross-validation gradient boosting machine learning models. Blood-related performances on these raw data subsets (cf. **Fig. 9**) were then compared to equivalent ML models built with scrambled metadata or shuffled samples. **a**, Comparison of blood-based ML models built on raw count data versus equivalent models built using scrambled metadata labels or shuffled count data using WGS samples from Harvard Medical School (HMS). **b**, Comparison of blood-based ML models built on raw count data versus equivalent models built using scrambled metadata labels or shuffled count data using WGS samples from Baylor College of Medicine (BCM). **c**, Comparison of blood-based ML models built on raw count data versus equivalent models built using scrambled metadata labels or shuffled count data using WGS samples from MD Anderson (MDA). **d**, Comparison of blood-based ML models built on raw count data versus equivalent models built using scrambled metadata labels or shuffled count data using WGS samples from Washington University (WashU). **e**, Comparison of blood-based ML models built on raw count data versus equivalent models built using scrambled metadata labels or shuffled count data using WGS samples from the Broad Institute. **a, b, c, d, e**, Pairwise two-sided Wilcox tests using performances from all ML folds, corrected for multiple hypothesis testing using the Benjamini-Hochberg method, are shown. ns: not significant (q>0.05); *: q≤0.05; **: q≤0.01; ***: q≤0.001; ****: q≤0.0001. BDN, blood derived normal.

We support the statement made by Gihawi *et al*., “to include only taxa with strong evidence of presence based on computational evidence, consideration of the likelihood of contamination and prior biological evidence that the taxa exist in the biological sample of interest.” Repeating our analyses would require external validation of taxa well-accepted to exist in various tumor types. To our knowledge, the independent (Weizmann, WIS), highly-decontaminated, multi-cancer cohort by Nejman *et al*.^2^ provides one such exemplary source of intra-tissue bacteria that have multiple levels of biological evidence for their existence. Thus, we subsetted our original Kraken data^1^ to WIS-overlapping genera (**Fig. 1a**) and repeated key analyses from our original manuscript; additionally, we have included several analyses without batch correction to demonstrate that the results do not critically depend on this factor, and we have included multiclass machine learning analyses to clearly show that mitigating the class imbalances still results in the same conclusion.

Comparing the original Kraken genera to the WIS-identified bacterial genera revealed substantial overlap, with 84.4% of WIS bacteria independently found by our TCGA analysis (**Fig. 1a**). Although virtually all versions of decontaminated data that we released captured the majority of WIS bacteria (FD, LCR, APCR, and PCCR datasets each identified >52% of them; **Fig. 1a**), the most stringently decontaminated feature set had substantially fewer (4.59% or just 10 WIS bacterial genera). These data fit the pattern we originally observed in our data, in which most typically expected commensals were discarded upon the most stringent filtering (Extended Data Fig. 6c-f in the original text). To proceed, we used the 184 WIS-overlapping bacterial genera present in our TCGA data to repeat batch correction using Voom and SNM, which were qualitatively and quantitatively evaluated using principal components analyses (**Fig. 1b**) and principal variance components analyses (**Fig. 1c**). Notably, despite using just 9.23% of the original Kraken-identified taxa (184/1993 genera), the post-batch corrected data again revealed that TCGA disease type explained more of the known variation than any other factor (**Fig. 1c**). These results suggested that it may be possible to discriminate between TCGA cancer types solely using WIS-overlapping bacterial abundances.

Thus, we implemented extensive one-cancer-type-versus-all-others ML analyses to quantitatively evaluate whether cancer types could be strongly distinguished using only this WIS-overlapping set, even though it contained <10% of our original features (**Fig. 2**). In order to evaluate the variances in model performances, we employed 10-fold cross validation, and by calculating the AUROC and AUPR on each of the 10 independent train-test splits, we estimated the concomitant 95% confidence intervals. These performance intervals were then compared to the known, null AUROC and AUPR values to see if overlap existed, as done extensively in our pan-cancer mycobiome manuscript^3^. Notably, all primary tumor-based ML models substantially exceeded their null performances (**Fig. 2a**), indicating that each cancer type was distinguishable solely using WIS-overlapping bacterial genera. Similarly, all tumor versus adjacent normal ML models showed better-than-null performances (**Fig. 2b**), and all blood-based ML models demonstrated the same conclusion (**Fig. 2c**). Collectively, even when using <10% of our original feature set, including those taxa identified by an independent, international group, all of our central conclusions are maintained.

Nonetheless, the primary tumor and blood analyses often contain imbalanced classes, sometimes to the extent of exhibiting one positive class instance for every hundred negative class examples. Therefore, we applied multiclass ML to test whether dozens of cancer types could be simultaneously discriminated solely using microbial nucleic acids. Performing this procedure among primary tumors indeed revealed very strong pan-cancer diagnostic performance (**Fig. 2d**). Directly addressing the concerns of Gihawi *et al*., the no information rate for the resultant multiclass confusion matrix was just 8.92%, and yet the raw accuracy was more than 6-fold greater (54.5%; mean balanced accuracy=75.99%) with a concomitant p-value <2.2×10^−16^; moreover, the average pairwise AUROC was 91.68%, much greater than the null expectation of 50%. Repeating the same procedure using all blood samples (**Fig. 2e**) showed even stronger simultaneous pan-cancer discrimination, again with a no-information rate of the resultant multiclass confusion matrix of just 8.63% but a raw accuracy more than 7-fold greater (66.07%, p<2.2×10^−16^; mean balanced accuracy=83.58%) and an average pairwise AUROC of 95.76% (vs. 50% for null models). These results strongly support the pan-cancer predictive value of tumor- and blood-based microbial nucleic acids, including when the analysis is restricted only to the WIS-overlapping bacterial genera.

Given the strength of these classifiers, we further sought to compare them to equivalent models with either scrambled metadata labels (e.g., cancer type) or shuffled count data. If the data were artificially distributed in such a way as to allow strong predictive performance in these circumstances, these analyses would reveal the unsuitability of the methods used—this method of control analyses was extensively undertaken in our pan-cancer mycobiome manuscript^3^. However, comparing the actual performance to ML models with scrambled metadata labels or shuffled data indeed revealed significantly worse performance, which corresponded with null expected values for primary tumor comparisons (**Fig. 3a**), tumor vs. adjacent normal comparisons (**Fig. 3b**), and blood comparisons (**Fig. 4a**). In fact, the only instance in this entire analysis that failed to show significance was tumor vs. adjacent normal tissue in esophageal cancer (ESCA, **Fig. 3b**), which was a type of cancer not included in the WIS cohort. These analyses therefore support the validity and strength of our original manuscript’s conclusions.

Because our original manuscript also evaluated predictive performances between cancer types when using blood samples from patients with early-stage (Ia-IIc) cancer, and differences between stage I and IV tumors, we repeated these analyses here using only the WIS-overlapping bacterial genera. Notably, the ability to distinguish between cancer types when solely using blood-derived microbial nucleic acids was equivalent to our original published results (**Fig. 4b**). Importantly, because WIS bacteria were only identified using tissue sections, this analysis reinforces the idea that a substantial source of blood-derived microbial nucleic acids is intratumoral bacteria, as we concluded in our mycobiome manuscript^3^. Examining differences between stage I and IV tumors revealed nearly identical performance for each cancer type to what we previously published (**Fig. 4c**). These data confirm and extend the findings stated in our original manuscript.

Each of these ML comparisons relied on using a batch-corrected version of the TCGA microbiome, which was necessary to address substantial technical variation in the data (**Fig. 1c**). In order to remove all doubt that the batch correction process artificially influenced the strength of the classifiers, we subset all raw data (WIS-overlapping features) to individual sequencing centers, experimental strategies (WGS or RNA-Seq), and a single sequencing platform (Illumina HiSeq, comprising 91.27% of samples), since these three variables accounted for most of the technical variation. We then repeated all of our ML approaches solely using raw data from these subsets, as done in our pan-cancer mycobiome manuscript^3^. This analysis revealed that, in all raw data subsets, primary tumors were always distinguishable using WIS-overlapping bacteria (**Fig. 5**). Moreover, implementing the same scrambled and shuffled ML control analyses revealed significantly better performance for the actual samples in every raw data subset (**Fig. 6**).

We continued this pattern of analysis for tumor vs. normal tissues using raw data subsets, finding that every model outperformed (**Fig. 7**) its null expected values, with the sole exception of lung adenocarcinoma at Harvard Medical School (**Fig. 7a**). Still, when aggregating cancer types in each sequencing center, the raw data subsets performed significantly better than matched scrambled or shuffled control models (**Fig. 8**). In the same manner, repeating the blood-related ML models using raw data subsets consistently showed strong performances that were always better than the null expected values (**Fig. 9**). This was complemented by these models also showing significantly better performances than scrambled or shuffled ML model counterparts (**Fig. 10**).

To conclude, all of our central conclusions of the original manuscript are replicated using a subset of <10% of the bacterial taxa obtained from an independent, tissue-centric cohort that escapes the contamination concerns raised in this paper. Our published pan-cancer mycobiome manuscript^3^ also affirms these findings using updated, state-of-the-art methods, some of which were adopted for use in this response. These additional analyses further substantiate confidence in the original findings of the manuscript, and were all possible using public data and code that we had already provided.

